# Single-cell multi-omic profiling of chromatin conformation and DNA methylome

**DOI:** 10.1101/503235

**Authors:** Dong-Sung Lee, Chongyuan Luo, Jingtian Zhou, Sahaana Chandran, Angeline Rivkin, Anna Bartlett, Joseph R. Nery, Conor Fitzpatrick, Carolyn O’Connor, Jesse R. Dixon, Joseph R. Ecker

## Abstract

Recent advances in the development of single cell epigenomic assays have facilitated the analysis of gene regulatory landscapes in complex biological systems. Methods for detection of single-cell epigenomic variation such as DNA methylation sequencing and ATAC-seq hold tremendous promise for delineating distinct cell types and identifying their critical cis-regulatory sequences. Emerging evidence has shown that in addition to cis-regulatory sequences, dynamic regulation of 3D chromatin conformation is a critical mechanism for the modulation of gene expression during development and disease. It remains unclear whether single-cell Chromatin Conformation Capture (3C) or Hi-C profiles are suitable for cell type identification and allow the reconstruction of cell-type specific chromatin conformation maps. To address these challenges, we have developed a multi-omic method single-nucleus methyl-3C sequencing (sn-m3C-seq) to profile chromatin conformation and DNA methylation from the same cell. We have shown that bulk m3C-seq and sn-m3C-seq accurately capture chromatin organization information and robustly separate mouse cell types. We have developed a fluorescent-activated nuclei sorting strategy based on DNA content that eliminates nuclei multiplets caused by crosslinking. The sn-m3C-seq method allows high-resolution cell-type classification using two orthogonal types of epigenomic information and the reconstruction of cell-type specific chromatin conformation maps.

## Introduction

Three-dimensional genome architecture is emerging as a critical feature of gene regulation in metazoan organisms ^1–3^. Chromatin conformation profiling using methods such as Hi-C provides experimental evidence of genomic features such as topologically associated domains (TADs) and enhancer-promoter interactions ^4–9^. Despite the increasing utility of these datasets, most existing chromatin conformation maps are generated from cell lines *in vitro* or from bulk tissues *in vivo ^4,8–11^*. While cell line data has enabled a greater understanding of the general principles of chromatin organization, it cannot fully represent the diversity of cell types that arise *in vivo*. Likewise, generation of chromatin interaction maps from bulk *in vivo* tissues precludes the analysis of cell-type specific chromatin organizations. Recent efforts using singlecell “omics” technologies aims to resolve this challenge by generating single-cell data from complex tissues that are then partitioned into the relevant distinct cell types *in silico* using data dimensionality reduction and clustering algorithms ^12–15^. Single-cell 3C or Hi-C therefore represent attractive strategies to resolve cell-type heterogeneity ^16–18^. However, current single-cell Hi-C profiles from cultured cells primarily capture cell cycle patterns ^16,19^. In this sense, it remains unclear whether single-cell Hi-C profiles will be suitable for partitioning into constituent cell types *in vivo*.

In contrast to single cell Hi-C data, single-cell DNA methylome datasets enable high-resolution cell-type classification, allowing the reconstruction of epigenomic maps from cell types in primary human tissues ^20, 21^. 3C or HiC methods capture chromatin configuration by performing proximity ligation with restriction digested genomic DNA in crosslinked nuclei ^22^. DNA methylation (mC) is fully preserved in the chimeric DNA molecules produced by 3C or HiC. Therefore it is feasible to jointly detect both long-range ligation junction and mC by analyzing DNA molecules generated by 3C or HiC using bisulfite sequencing. When applied to single-cell analysis, cell type identification using both chromatin conformation and mC signatures could provide superior resolution than using either feature alone. Joint analysis of chromatin conformation and mC can also facilitate the study of cross-talk between the two epigenomic features.

Here we describe a method, single-nucleus methyl-3C sequencing (sn-m3C-seq), to jointly profile chromatin conformation and DNA methylation from the same cell. Bulk and single-cell m3C-seq profiles accurately recapitulate chromatin architectures of mouse embryonic stem cells (mESCs). Using a mixed species design, we show that current *in-situ* single-cell 3C-seq or Hi-C method produce a significant amount of nuclei multiplets due to inter-nuclei crosslinking. By stringently sorting particles with 2n DNA content, we are able to eliminate nuclei multiplets during fluorescent-activated nuclei sorting (FANS). Finally, we show that sc-m3C-seq can allow for the unbiased analysis of distinct mouse cell types.

## Results

### Joint profiling of chromatin conformation and DNA methylation from the same DNA molecule

Here we describe the development of a novel method for joint profiling of 3D chromatin structure and mC that we term *in situ* 3C followed by mC analysis by sequencing (m3C-seq). The outline of this method is described in Fig.1. First, we perform restriction enzyme digestion and ligation on fixed nuclei, as is typically performed in an *in situ* 3C experiment ^8,22^. When performed as a bulk assay, the ligated 3C nuclei are subject to proteinase digestion and bisulfite conversion, and libraries are constructed similar to our previous snmC-seq2 method using a random primed DNA synthesis and addition of downstream adaptor using Adaptase^20,23^. The procedure is similar for single-nucleus reactions, except that single nuclei are dispensed into 384 well PCR plates using FANS (Fig.1).

**Figure 1.**
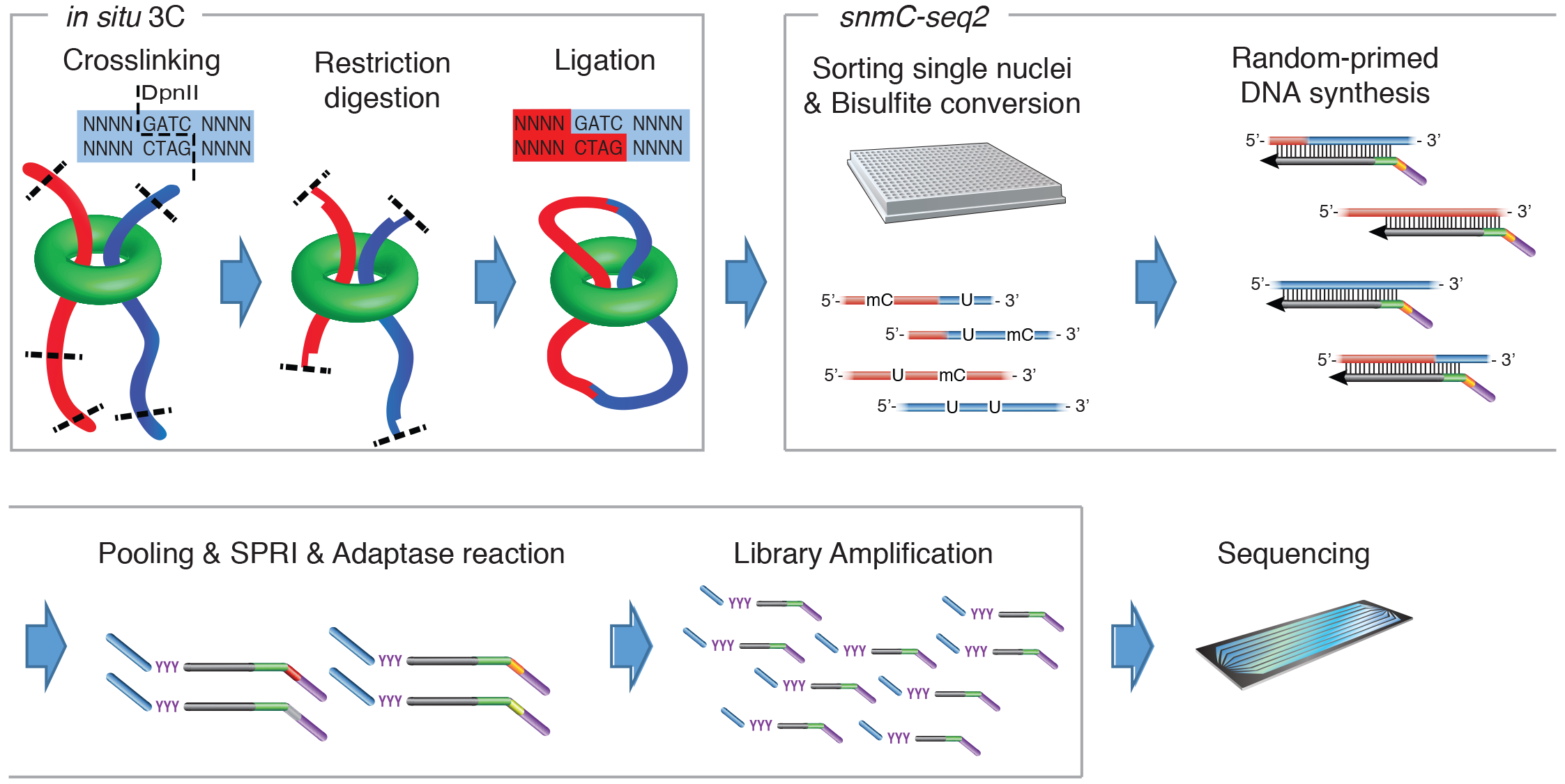
Outline of the single-nucleus methyl-3C sequencing (sn-m3C-seq) method. Samples are crosslinked with formaldehyde in a typical in situ 3C/Hi-C experiment. The nuclei are isolated and permeabilized. Chromatin is then digested with a restriction enzyme of interest and ligated to preferentially join regions that were originally in close 3D space. Importantly, the digestion and ligation reactions are occurring inside intact nuclei. Nuclei are then sorted into 384 well PCR plates and subjected to bisulfite conversion. Single-cell DNA methylome libraries were prepared using snmC-seq2 method. Briefly, DNA in each well is then labeled by random primers which contain a specific barcode. After pooling of wells with unique random primer barcodes, the 3’-end of random-primed DNA synthesis products are then modified using an Adaptase reaction to attach a 3’-adaptor.

To evaluate the consistency of chromatin contact maps generated by m3C-seq, we first performed bulk m3C-seq experiments for mESC in parallel with conventional bulk *in situ* 3C and Hi-C sequencing experiments. Both Hi-C/3C and bisulfite conversion can present challenges for proper read alignment. In the case of Hi-C, this is due to the presence of chimeric reads where the read covers the ligation junction site. In the case of bisulfite treated sequencing libraries, this is due to the conversion of unmethylated cytosines to uracils, which are read as thymines during sequencing. In order to ensure accurate alignment of the m3C-seq data, we developed TAURUS-MH (**T**wo-step **A**lignment with **U**nmapped **R**eads **U**sing read **S**plitting for **M**ethyl-**H**iC), a mapping pipeline for m3C-seq data using a hybrid of ungapped and manual read splitting alignments (Fig. 2a). Sequencing reads were first mapped to a *in silico* bisulfite converted genome using Bismark calling an ungapped aligner (bowtie1)^24^. Unmapped reads were further processed using a read-splitting procedure, which split reads into 3 segments followed by ungapped mapping of the split reads. To evaluate the performance of our pipeline, we compared TAURUS-MH to a previously described pipeline using gapped alignment (BWA-MEM) using a reference Hi-C dataset ^25^. We also compared our pipeline with BWA-METH^26^, which is designed for bisulfite sequencing data using BWA-MEM. The comparison with BWA-METH was performed using simulated bisulfite converted Hi-C reads. When compared with BWA-METH, our pipeline showed 19.43% higher in mappability (86.12% vs. 66.69%, Fig. 2b), 3.64% higher in accuracy (97.86% vs. 94.22%, Fig. 2c), and 13.41% higher long-range *cis* contacts (42.79% vs. 29.38% from total fragments and 49.68% vs. 44.06% from mapped fragments, Fig. 2d). In summary, our pipeline showed improved performance both in accuracy and sensitivity, compared with current methods.

**Figure 2.**
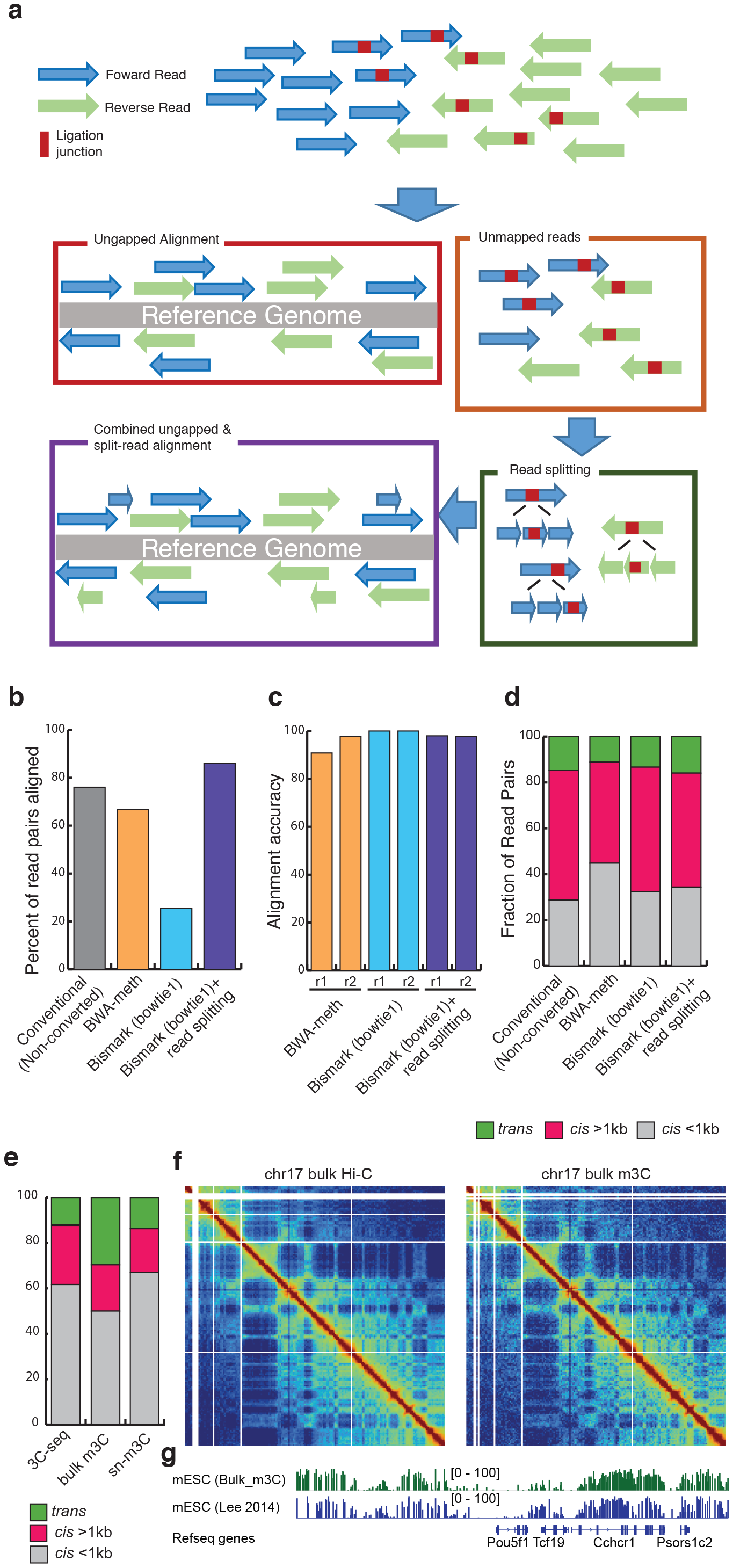
Data processing and analysis of m3C-seq sequencing reads. **(A)** Overview of 3C-seq mapping. Reads are aligned using Bismark calling ungapped aligner bowtie1. To rescue reads that did not align due to the presence of a ligation junction within the reads, we split unmapped reads into 3 equal segments and these are realigned. Successfully aligned reads are then manually paired, deduplicated, and then processed for DNA methylation and chromatin contact profiles. **(B)** Percent of aligned reads as a pair. Reads derived from non-bisulfite treated regular Hi-C sequencing are converted C to T (read1) and G to a (read2) in silico and aligned using BWA-meth, Bismark (bowtie1), and Bismark (bowtie1) followed by split-read alignment. Alignment of non-converted reads using conventional alignment pipeline is used as a standard. **(C)** Alignment accuracy of different alignment strategies compared with conventional Hi-C alignment using *in silico* converted reads. **(D)** Fraction of read pairs with cis short-range reads (cis < 1kb), cis long-range interactions (cis > 1kb), and trans interactions (trans) using different alignment strategies. **(E)** Similar to panel (D), but showing cis short-range reads (cis < 1kb), cis long-range interactions (cis > 1kb), and trans interactions (trans) for actual 3C-seq (without conversion), bulk m3C-seq (with conversion, from the same sample as bulk 3C-seq), and single-nucleus m3C-seq results. **(F)** Contact maps from chromosome 17 for conventional bulk Hi-C and bulk m3C data. **(G)** DNA-methylation profiles near the Pou5f1 gene for conventional bulk MethylC-seq as well bulk m3C-seq.

Having developing a robust mapping pipeline, we then analyzed chromatin contacts from libraries generated from bulk m3C-seq with a matched 3C-seq library. Bulk m3C-seq libraries showed a comparable fraction of long-range (>1kb) intra-chromosomal ligation events compared to the control 3C-seq library (26.6% in 3C-seq and 19.0% in m3C-seq) (Fig. 2e). Surprisingly, we observed much more frequent inter-chromosomal ligation events in m3C-seq than in 3C-seq libraries (12.24% in 3C-seq and 30.0% in m3C-seq) (Fig. 2e). Since snmC-seq2 is based upon random-primed DNA synthesis ^23^, we speculate that the inter-chromosomal ligation is an artifact caused by spurious hybridization and polymerase extension between products of random-primed DNA synthesis. Consistent with this conjecture, no inter-chromosomal ligation events were found if we use reads that are aligned within 200bp of DpnII restriction sites, supporting the hypothesis that they are generated by a different mechanism from restriction digestion and ligation. We further hypothesized that spurious inter-chromosomal ligation is dependent on DNA concentration, which is positively correlated with the frequency of intermolecular interaction. Therefore, we expected sn-m3C-seq data to show reduced inter-chromosomal ligation since the random-primed DNA synthesis reaction for sn-m3C-seq contains a much lower DNA concentration. Indeed, we found a background level (15.11%) of inter-chromosomal ligation in sn-m3C-seq datasets, which is similar to what was observed in a regular 3C-seq experiment (12.24%). Finally, we compared the contact maps and DNA methylation profiles generated by our bulk m3C-seq with conventional Hi-C and MethylC-seq data generated from mESC (Fig. 2f,g)^27^. We observed strong agreement in the contact maps (Fig. 2f). Consistent with visual examination, the matrices showed high correlation with bulk Hi-C data as measured using the HiCrep tool (Pearson correlation = 0.979) ^28^. HiCrep is a tool designed to compare Hi-C datasets using stratum adjusted correlation coefficients in order to account for unique features of Hi-C datasets that can lead to artificially high correlation coefficients when using conventional metrics like Pearson or Spearman correlation. Similarly, we observed strong concordance of methylation profiles from bulk m3C-seq with existing MethylC-seq datasets for mESC (Fig. 2g, Pearson correlation = 0.977).

### Fluorescent-activated nuclei sorting by DNA content excludes nuclei multiplets caused by inter-nuclei crosslinking

To generated sn-m3C-seq profiles, fluorescent-activated nuclei sorting (FANS) was applied to *in situ* 3C treated nuclei preparation to dispense single nuclei into 384 well PCR plates followed by snmC-seq2 single-cell methylome library preparation. We speculated that the crosslinking step of 3C or HiC may lead to the crosslinking of randomly interacting nuclei and cause more frequent nuclei multiplets compared to regular single-cell methylome preparation. In order to quantify the frequency of nuclei multiplets, we designed a mixed species experiment to crosslink an mixture of mESC and human GM12878 nuclei (Table S1). Consistent with our speculation, we found evidence of nuclei multiplets in 22.8% of PCR plate wells, using a threshold that requires more than 10% of reads mapped to both mouse and human genomes (Fig. 3a). This is despite performing strict singlet gating using forward scatter and trigger pulse width. We found that the species multiplets were due to formaldehyde crosslinking, as experiments where mouse and human cells were crosslinked separately and then combined for the subsequent experimental steps eliminated wells showing reads mapped to both species (Fig. 3b), supporting our hypothesis that nuclei multiplets are caused by inter-nuclei crosslinking instead of FANS steps or sequencing barcode crossover.

**Figure 3.**
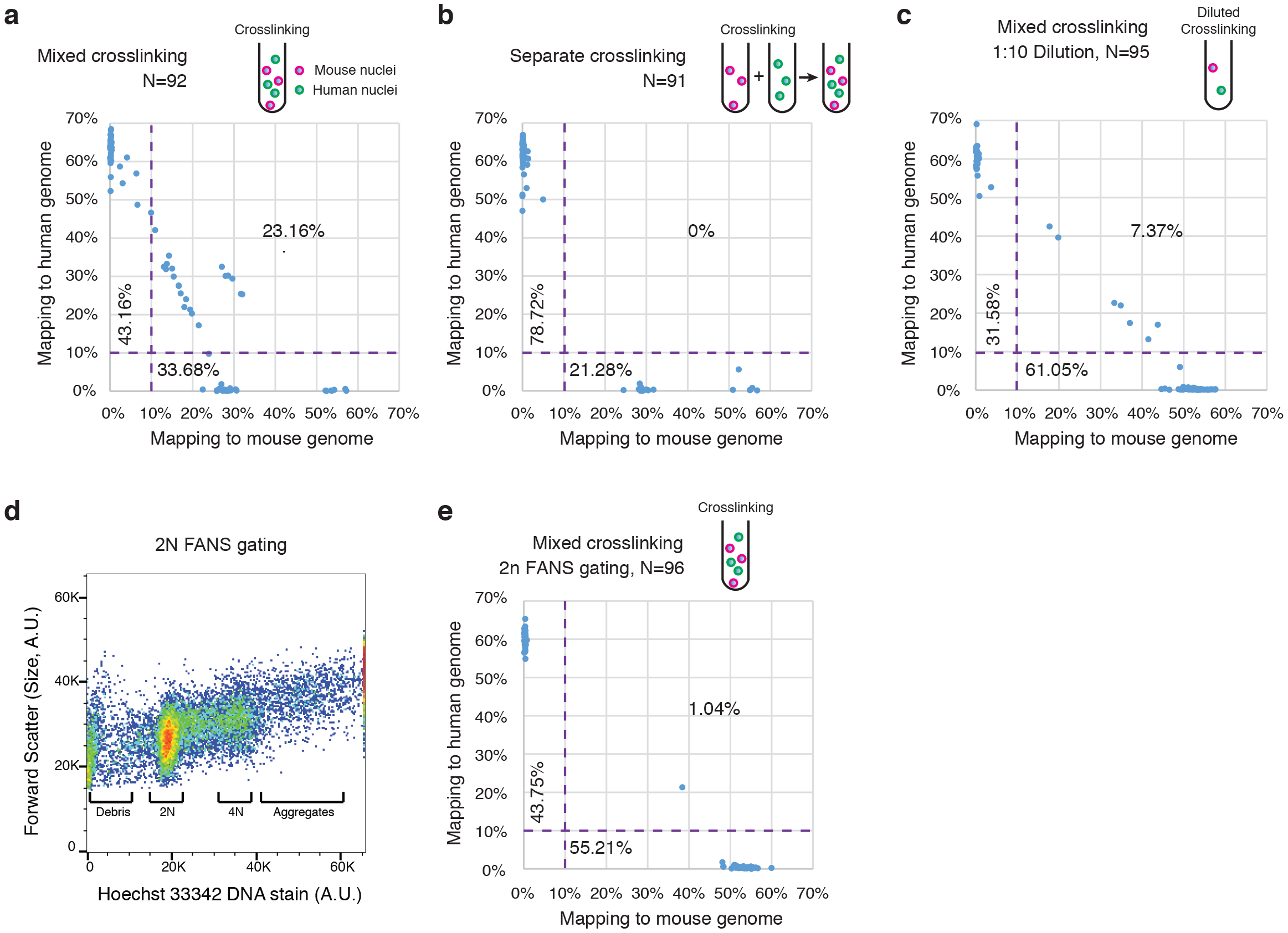
FANS by DNA content excludes nuclei multiplets. **(A)** Single-nuclei FANS following standard *in situ* 3C procedure using an mixture of mESC and GM12878 results in a high fraction of wells containing both mouse and human nuclei. **(B)** Separate crosslinking of mESC and GM12878 nuclei followed by pooling and FANS eliminated wells containing both mouse and human nuclei. **(C)** Crosslinking under diluted condition reduced nuclei multiplets. **(D)** FANS selecting for 2N genomic content. **(E)** FANS selecting for 2N genomic content excluded the vast majority of nuclei multiplets.

To reduce the formation of nuclei multiplets, we first tested a strategy in which crosslinking is carried out with a nuclei preparation diluted 10-folds, potentially reducing random interactions between nuclei. Performing crosslinking at a diluted condition substantially reduced the frequency of nuclei multiplets to 7.4% of the wells compared to 22.8% of wells for the undiluted condition (Fig. 3c). We also tested a strategy in which sort nuclei are stringently selected during FANS with a 2n genomic DNA content to exclude particles containing more than one nucleus (Fig. 3d). Applying this strategy to the mixed species design effectively excluded nuclei multiplets with only 1% of wells showing reads mapped to both mouse and human genomes (Fig. 3e). Thus we concluded that FANS with stringent gating for 2n DNA content is the most effective approach for excluding nuclei multiplets from downstream single-nuclei analysis.

### Bulk m3C-seq and sn-m3C-seq profiles recapitulate chromatin conformation contact maps

We compared chromatin contacts and DNA methylation patterns generated from sn-m3C-seq with that from bulk Hi-C, m3C-seq, and bulk MethylC-seq. We generated 192 mouse ES cells sn-m3C-seq DNA methylation and chromatin conformation profiles (Table S1). The data from the 192 single cells was then merged to reconstruct bulk contact maps and methylation profiles. The merged sn-m3C-seq contact maps were highly consistent with contact maps from conventional Hi-C and from bulk m3C-seq experiments (Fig. 4a,b). We quantitatively assessed the similarity of the sn-m3C-seq and Hi-C contact maps using HiCrep ^28^. We compared our pooled sn-m3C-seq and Hi-C maps with existing public mouse ES cell Hi-C data. In addition, we used bulk Hi-C data recently generated from Bonev et al. from neural progenitor cells (NPC) and cortical neurons (CN) to act as an outgroup for comparison ^9^. Performing hierarchical clustering on a matrix of correlation coefficients generated by HiCrep, we observed that our sn-m3C-seq clustered with previous Hi-C data from mouse ES cells as well was with Hi-C data generated as part of this study, while the CN and NPC datasets clustered separately (Fig. 4c). In addition, we were able to observe ES cell specific contacts that were consistent between pooled sn-m3C-seq and Hi-C data, such as the ES cell specific enhancer-promoter contacts at the Sox2 locus (Fig. 4c). Lastly, we also compared the methylation profiles generated in the m3C-seq and sn-m3C-seq with existing MethylC-seq profiles in mouse ES cells, NPCs, CNs, and frontal cortex datasets. We observed a high correlation of the methylation profiles generated by pooled sn-m3C-seq with previously generated bulk MethylC-seq experiments (Pearson correlation = 0.89, sn-m3C-seq vs. mESC Lee 2014) (Fig. 4d) ^27^. We were also able to observe characteristic hypomethylated regions in the promoters of cell-type specific genes, such as the pluripotency genes Dppa4 and Dppa2 (Fig. 4d). In summary, these results demonstrate that the contact maps and methylation profiles generated by sn-m3C-seq and bulk m3C-seq are consistent with published Hi-C and MethylC-seq data.

**Figure 4.**
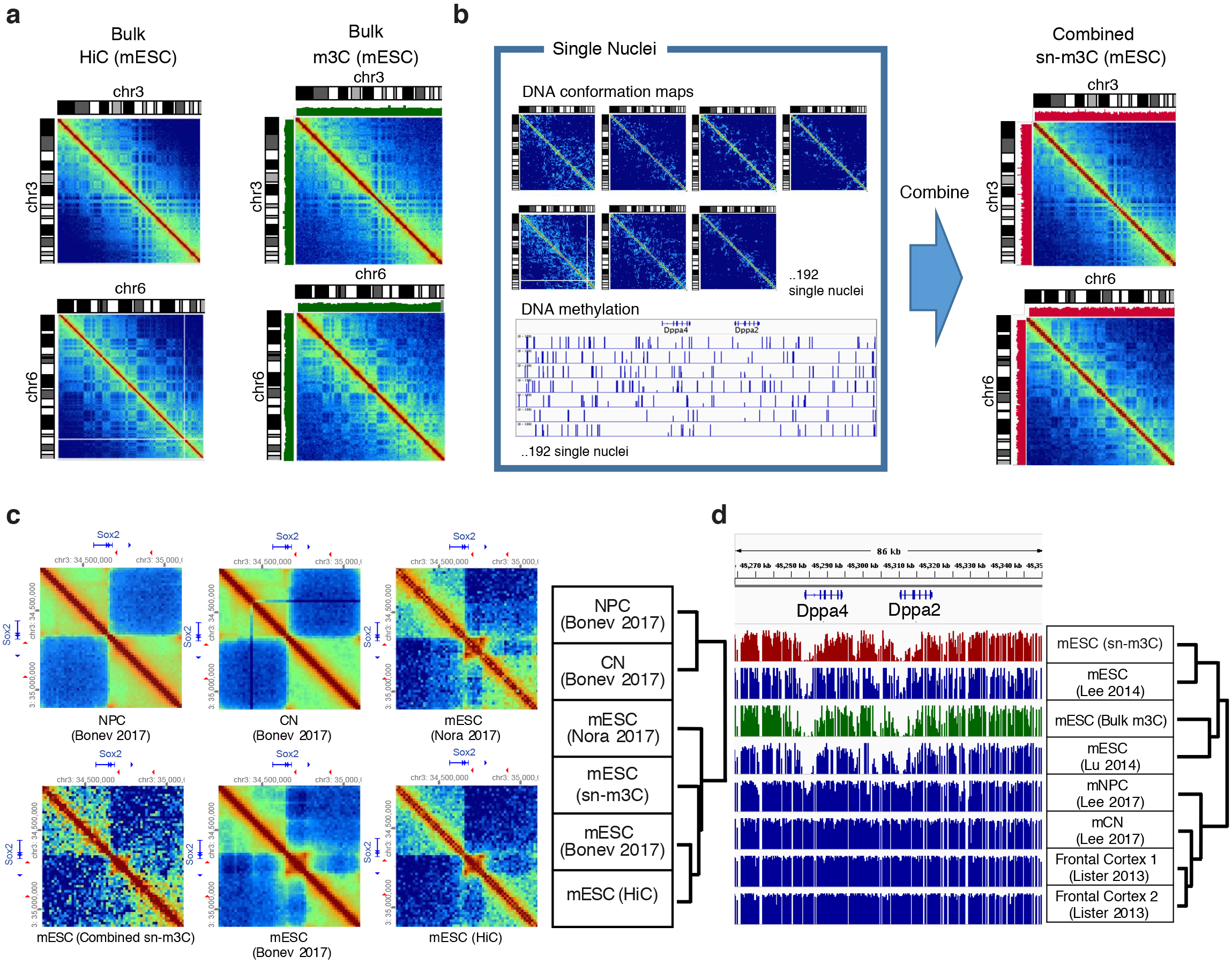
Bulk and single-nucleus m3C-seq recapitulate the chromatin architecture of mouse embryonic stem cells. **(A)** HiC and bulk m3C-seq profiles show consistent chromosome level chromatin architectures. Green bar plot shows CpG methylation level from m3C-seq. **(B)** Reconstructed mESC chromatin conformation map from sn-m3C-seq profiles is highly consistent with the maps generates using Hi-C or bulk m3C-seq. Red bar plot shows CpG methylation level from sn-m3C-seq. **(C)** Bulk and single-nucleus m3C-seq identify the mESC specific chromatin conformation surrounding Sox2 locus. **(D)** Bulk and single-nucleus m3C-seq recapitulate the mESC specific depletion of DNA methylation at Dppa2/4 locus.

### sn-m3C-seq profiles separate mouse cell types

To test the feasibility of cell type identification using sn-m3C-seq profiles, we analyzed sc-m3C-seq data from mESCs and a mouse mammary gland cell line, NMuMG. We sorted 192 cells for each sample and processed the samples for sn-m3C-seq, and sequenced the libraries to obtain on average 1.4 million read pairs per cell for mESC, and 1.7 million reads pairs for NMuMG (Table S1). After filtering for cells containing at least 100,000 uniquely mapped reads, our dataset consisted of 152 mouse mESCs and 96 mouse mammary gland NMuMG cells. 68.77% of mESC and 66.49% of NMuMG read pairs can be uniquely mapped to mouse genome. The mESC sn-m3C-seq dataset contains on average 1,229,602 (standard error s.e.=126,534) uniquely mapped reads per cell, with 15% (s.e=0.31%) of reads indicating long-range (>1kb) chromatin interactions. The NMuMG dataset contains on average 1,433,470 (s.e.=125,560) uniquely mapped reads per cell, with on average 8.64% (s.e.=0.35%) of reads indicating long-range (>1kb) chromatin interactions.

To partition cells into separate cell types, we used CpG methylation patterns from single cells, analogous to what we have done previously in human and mouse neurons ^20^. We binned the genome into 100kb bins and calculated the average CpG methylation level in each bin for each cell. Principal component analysis (PCA) of the resulting single cell CpG methylation patterns resulted in two clear clusters, one corresponding of NMuMG cells and the other nearly entirely composed of ES cells (Fig. 5a). Having partitioned the single cells into constituent clusters, we then pooled the sc-m3C-seq data from ES and NMuMG cells and analyzed the chromatin contact maps for differences. We were able to identify chromosomes in each lineage with distinct A/B compartment signatures between the two cell types (Fig. 5b) as well as identifying local differences in Hi-C contacts (arrows in Fig. 5c,d).

**Figure 5.**
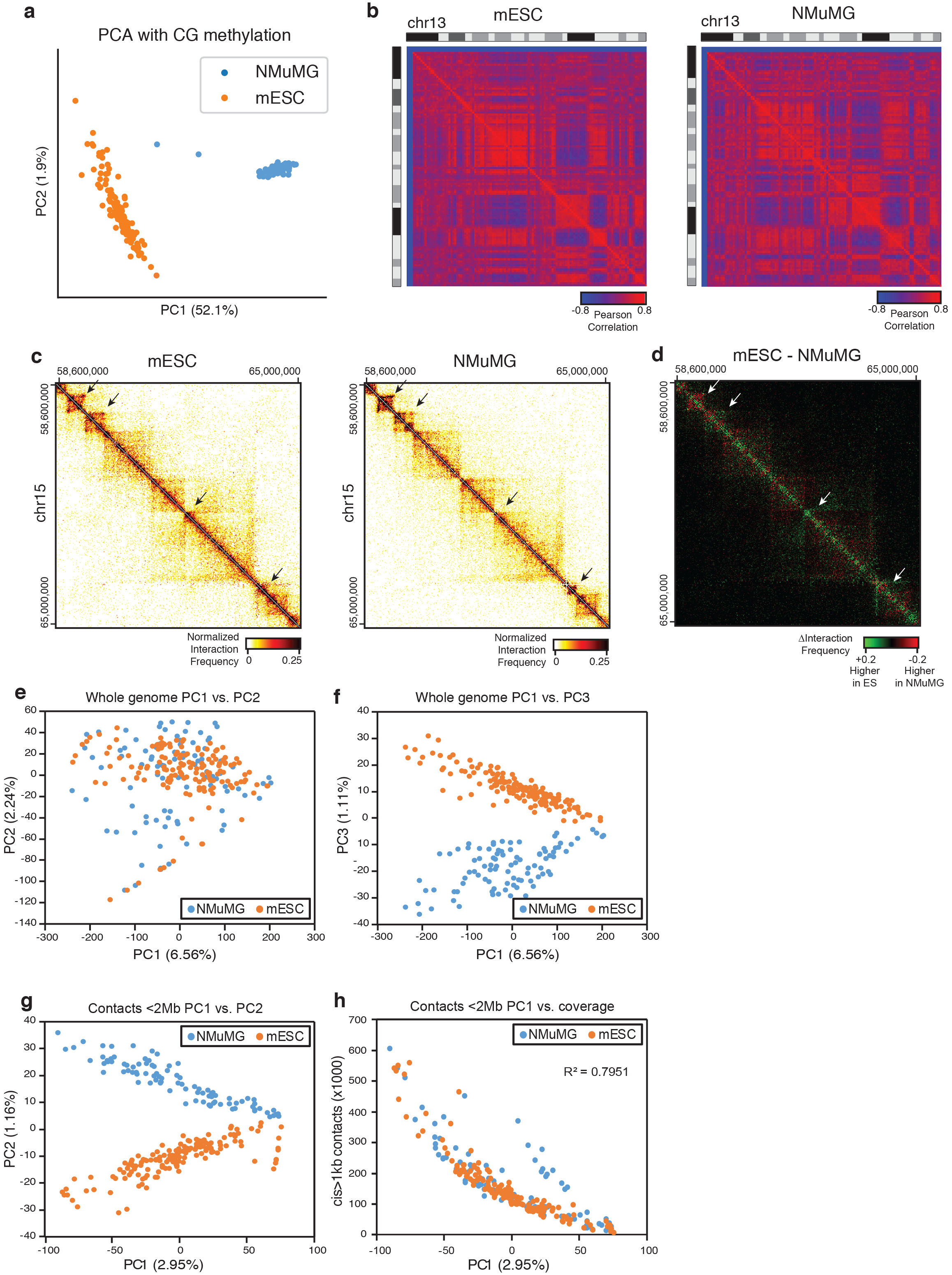
Single-nucleus m3C-seq reconstructs cell-type specific chromatin conformation maps. **(A)** Principal component analysis of single cell DNA-methylation profiles of mouse ES cells and NMuMG cells reliably distinguishes the two cell populations. **(B)** Chromosome wide Pearson correlation matrix from pooled sc-m3C-seq maps for ES cells and NMuMG cells demonstrates cell type specific patterns, corresponding to cell type specific A/B compartment signals. **(C)** Contact profiles from sc-m3C-seq data in a 6.4Mb stretch of chromosome 15 show cell type specific contacts in both ES and NMuMg cells. **(D)** Heat map of differential interaction frequencies between mESC and NMuMG cells shown in panel C. Regions in green are stronger in ES cells, regions in red are stronger in NMuMG. **(E)** Principal component analysis of whole genome contact matrices from sc-m3C-seq fails to distinguish ES from NMuMG cells using PC1 and PC2. (POV: Percentage of variance) **(F)** Principal component analysis of whole genome contact matrices from sc-m3C-seq distinguishes ES from NMuMG cells using PC3. **(G)** Principal component analysis of local interactions (<2Mb) sc-m3C-seq distinguishes ES from NMuMG cells using PC1 and PC2. **(H)** Correlation of PC1 and per cell contacts demonstrates that PC1 largely corresponds to coverage.

We also analyzed the ability to partition sc-m3C-seq into the relevant cell types using DNA-methylation patterns compared with Hi-C based contacts. As stated previously, DNA-methylation profiles could easily distinguish between ES and NMuMG cells using the first principal component (PC) alone, which explains 52.1% of total variance (Fig. 5a). In contrast, PCA of genome wide Hi-C contacts could not distinguish between ES and NMuMG cells using the first two PCs (Fig. 5e), but the third PC did clearly separate these two cell types (Fig. 5f). We also carried out PCA using Hi-C based contacts separated by less than 2 Mb in the genome, as we reasoned that the matrices were less sparse for these local interactions. In this case, we were able to distinguish the two cell types using the first two PCs (Fig. 5g), where the ability to distinguish the two cell types was determined by the second PC. We also observed that the first PC was highly correlated with the per cell sequence coverage (Fig. 5h), such that the samples with higher coverage were more reliably separated by the second PC. This suggest that using PCA based cell type identification using Hi-C contacts is highly dependent on per cell sequence coverage. In contrast, DNA methylation profiles easily distinguish between the two cell types using PC1 values alone (Fig. 5a), indicating the robustness of cell type classification from DNA-methylation profiles. These results underscore the importance of jointly profiling DNA-methylation along with chromatin conformation to reliably distinguish cell types in single cell experiments.

## Discussion

Cell-type specific chromatin conformation maps can potentially provide a valuable addition to other single cell modalities for the creation of cell type atlases ^29^. This information complements single-cell transcriptomes and the annotation of regulatory elements using single-cell epigenomic profiles, to provide a more comprehensive description of gene regulatory activities. However, it is currently unclear how well singlecell Hi-C/3C methods alone can distinguish unique cell-types in a heterogenous population. To enhance the cell-type signature in single-cell chromatin conformation data, we devised a method to allow jointly profiling of chromatin interaction and DNA methylation from a single nucleus. Consistent with previous single-cell methylome studies, sn-m3C-seq allows unequivocal clustering of cell types, which can then guide the reconstruction of high-quality cell-type specific chromatin conformation maps.

Our results indicate that single cell contact profiles alone can distinguish between drastically different cell types such as mESC and NMuMG. However, the confidence of cell type separation is highly dependent on sample coverage and downstream processing. It is unclear how well these contact map may be able to distinguish between highly related cell types (such as different sub-types of neurons) present in complex tissue samples. In contrast, single-cell DNA methylation profiles can easily distinguish constituent cell types.by examining DNA methylation profiles associated with various well-known marker genes/loci.

Our study also made a critical improvement to current single-cell Hi-C/3C methods by excluding nuclei multiplets using FANS with 2n DNA gating strategy. Cell or nuclei multiplets are a major source of noise in single-cell studies and can lead to the identification of artifactual cell populations. Our modified FANS method will enable the generation of high quality single cell chromatin conformation maps. Similar to snmC-seq2, sn-m3C-seq is compatible with high throughput approaches, allowing analysis of thousands of single nuclei in a single experiment. Sn-m3C-seq is thus well suited for large-scale cell atlas projects and can contribute to a comprehensive analysis of gene regulatory activity with cell-type specificity.

## Data availability

Raw data and processed data are available from NCBI GEO accession GSE124391.

## Code availability

The source code used is publicly available at https://github.com/dixonlab/Taurus-MH

## Acknowledgements

This work was supported by NIH grant 5R21HG009274 to J.R.E and DP50D023071 to J.R.D. J.R.E is a Howard Hughes Medical Institute investigator. J.R.D is also supported by the Leona M. and Harry B. Helmsley Charitable Trust grant No. 2017-PG-MED001 and a grant from the Salk Institute Innovation Research Fund. This work was also supported by the Flow Cytometry Core Facility of the Salk Institute with funding from NIH-NCI CCSG: P30 014195.

## Material and Methods

### Cell culture

Mouse ES cells (E14TG2a) were purchased from American Type Culture Collection (ATCC CRL-1821). ES cells were grown in DMEM media (Corning 10-013-CV) supplemented with 10% HyClone FBS (Fisher SH3007003E), 1X MEM Non-essential amino acids (ThermoFisher 11140050), 1X Glutamax supplement (ThermoFisher 35050061), 1X ß-mercaptoethanol (Millipore ES-007-E), 100U/mL Penicillin-Streptomycin (ThermoFisher 15140122), and 1000U/mL Leukemia Inhibitory Factor (Millipore ESG1107). ES cells were cultured in feeder free conditions on 0.5% gelatin coated plates.

GM12878 cells were obtained from Coriell Institute for Medical Research. GM12878 cells were grown in RPMI-1640 medium (ThermoFisher 11875093) supplemented with 15% Fetal Bovine Serum (Corning 35-010-CV) and 100U/mL Penicillin-Streptomycin (ThermoFisher 15140122).

NMuMg cells (RBRC-RCB2868) were obtained from the RIKEN BioResource Center. NMuMg cells were grown in DMEM (Corning 10-013-CV) with 10% Fetal Bovine Serum (Corning 35-010-CV), 10**μ**g/mL Insulin (ThermoFisher 12585014), and 100U/mL Penicillin-Streptomycin (ThermoFisher 15140122).

All cell lines were routinely tested for mycoplasma contamination and tested negative.

### *in situ* Hi-C and 3C

*in situ* Hi-C was performed as previously described using the MboI restriction enzyme ^8^. *in situ* 3C experiments were performed based on the *in situ* Hi-C protocol with minor modifications. Briefly, prior to fixation, adherent cells were trypsinized, counted, and collected by centrifugation, and suspension cells were counted and collected by centrifugation. Cells were resuspended in culture media at a concentration of 1×10^6^ cells per mL of media and fixed in 1% formaldehyde for 10 minutes at room temperature with shaking. For standard species mixture experiments, equal numbers of mouse and human cells were combined into a single tube prior to fixation. For the 1:10 dilution species mixture experiment, cells were resuspended at a concentration of 1×10^5^ cells per mL of media prior to fixation. For the species mixture experiments where samples were mixed after fixation, each cell type was fixed independently as described above and combined at later stages in the protocol. *in situ* Hi-C samples were digested with the MboI restriction enzyme and processed as described previously. For *in situ* 3C experiments, samples were digested with Dpn 11 enzyme overnight at 37ºC with gentle mixing. The following day, the sample was incubated at 62°C for 10 minutes to inactivate the restriction enzyme. The typical biotin fill in step in the Hi-C protocol is omitted. The sample is then ligated for 4 hours at room temperature with T4 DNA ligase in the same manner as in *in situ* Hi-C experiments. The sample is then stained with Hoechst (0.1μg/μL) for the final 30 minutes of the ligation step. The sample is then passed through a 40 μM nylon cell strainer (Corning 431750) into a FACS tube prior to sorting. As a quality control step, 10% of the sample is taken for conventional library preparation and sequenced using shallow sequencing on a MiSeq.

### Fluorescent-activated nuclei sorting (FANS)

FANS was performed at the Salk Institute Flow Cytometry Core Facility using a BD Influx cell sorter. A 100 micron nozzle tip was used, with 1 x PBS as sheath fluid (sheath pressure was set to 18.5PSI) with sample and collection cooling set to 4 degrees. The gating strategy for selecting intact, single, Hoechst labelled nuclei from debris was as follows: nuclei were first gated based on Forward Scatter (FSC) and Side Scatter (SSC) pulse height, then multiplet exclusion gating was applied (forward scatter and side scatter pulse width). Finally, nuclei of specific DNA content were selected (e.g. 2N) by virtue of Hoechst fluorescence intensity. Individual nuclei were deposited into wells of 384-well plate using the Single Cell (1-drop) mode. In preparation for 384-well plate deposition, 20-30 particles (e.g. calibration beads) were sorted onto a transparent plastic plate cover for alignment calibration. 20-30 particles are then directly sorted into the wells for final visual confirmation of alignment precision.

### Bulk and single-cell methylome library preparation

Libraries for bulk and single-cell methylomes were generated using snmC-seq2. A detailed step-by-step bench protocol for snmC-seq2 is provided as Supplement Methods in Luo et al. (2018) ^23^. Bulk methylome libraries were prepared manually using individual tubes. Single-cell methylome libraries were prepared using a Tecan Evo 100 robotic platform as described in Luo et al. (2018) ^23^. Sequencing libraries were sequenced using Illumina MiSeq and HiSeq 4000 instruments in PE150 mode.

### Data Processing

Raw reads were trimmed first 25bp and last 3bp of both read1 and read2 to remove random primer sequence and adaptase low complexity tails. For the alignment, Bismark is used^24^. C to T converted and G to A converted reference genomes are prepared for mm10 and hg38 using bismark_genome_preparation. Each read end is mapped separately using Bismark with Bowtie1 with read1 as complementary (always G to A converted) and read2 (always C to T converted) as original strand. After first alignment, unmapped reads are retained and splitted into 3 pieces by 40bp, 32bp, and 40bp after removing 5bp of both ends results in having 6 reads (read1 and read2). Six reads derived from unmapped reads are mapped separately using Bismark Bowtie1. All aligned reads are merged into BAM using picard that is query name sorted. The fragments with all the mapped reads aligned to the same positions are considered as duplicates and removed and DNA methylation tracks are generated. For each fragment, the outermost aligned reads are chosen for the chromatin conformation map generation.

### Cell type identification using DNA methylation signature

CG methylation levels (mCG) are computed for every non-overlapping 100kb bins across the genome in each single cell. The bins with more than 20 CG basecalls in more than 95% of cells were selected for further analysis. Bin-level mCG levels were normalized by global mCG of each cell. We imputed the mCG in each bin with less than 20 CG basecalls by using the mean mCG of that bin across all the cells having more than 20 CG basecalls in the bin.

### Cell type identification using 3D genome structure

We generated contact map using 100kb bin in each cell (152 mESC and 96 NMuMG). The interaction frequency of each bin is normalized by dividing average interaction frequency of the bins at the same distance interactions. The bins that are covered with more than 100 cells were filtered (n=19357) and used for first PCA analysis (Fig. 5e,f). The bins with interaction distance of less than 2Mb are filtered and used for the second PCA analysis (n=18004)(Fig. 5g,h).

